# Tsallis-Gated Autoencoder: A Nonextensive Physics-Informed Approach for Unsupervised Anomaly Detection in Glioblastoma Multiforme RNA-seq Data

**DOI:** 10.64898/2026.05.13.724767

**Authors:** Sergio Assuncao Monteiro, Fabricio Alves Barbosa da Silva

## Abstract

Glioblastoma multiforme (GBM) is characterised by profound genomic heterogeneity and heavy-tailed gene-expression distributions that challenge conventional machine-learning methods. We introduce the *Tsallis-Gated Autoencoder* (Tsallis-GAE), a physics-informed architecture that replaces classical softmax attention with a learnable Tsallis *q*-softmax followed by mean-field smoothing iterations, motivated by recent work on curved statistical manifolds and dense associative networks. Trained on the full TCGA-GBM RNA-seq cohort (391 samples, top 2,000 high-variance genes) under a rigorous 80/20 hold-out protocol, the Tsallis-GAE achieves a mean AUC-ROC of **0.977** ± **0.002** across five independent seeds, compared to 0.906 ± 0.003 for a matched-capacity Vanilla autoencoder trained under the identical protocol. The matched-capacity Vanilla autoencoder is statistically indistinguishable from a LocalOutlierFactor baseline (AUC 0.906 vs 0.906), confirming that the +0.07 AUC gain over the Vanilla AE stems from the gated attention architecture rather than from the use of a neural network per se. A fixed-*q* Softmax-AE ablation (*q* ≡1 by construction) achieves AUC 0.976 ±0.001, only +0.001 below the Tsallis-GAE (DeLong *p* = 0.44); the physically meaningful contribution of the learnable *q* is its spontaneous convergence to the non-extensive regime described below. The three attention blocks each carry an independent learnable entropic index *q*; across 5 seeds ×3 blocks = 15 measurements, *q* converges spontaneously to 1.554± 0.019, strictly bounded away from the Boltzmann–Gibbs limit *q* = 1 and in the moderate non-extensivity regime characteristic of complex biological systems. Cross-detector validation against OneClassSVM and LocalOutlierFactor pseudo-labels yields Tsallis-GAE AUCs of 0.998 and 0.992 respectively, indicating that the learned representation captures anomaly structure intrinsic to the data rather than the decision boundary of any single labeling heuristic. We declare that DeLong’s paired test on the present test-set size (*n* = 79) does not certify the +0.07 AUC gap as formally significant (*p*≈ 0.26); a 5-fold cross-validation over the full cohort, which would supply the needed statistical power, is left to future work. The source code is available upon reasonable request to the corresponding author.

## 1 Introduction

Glioblastoma multiforme (GBM) remains the most aggressive primary brain tumor in adults, with a median survival of only 12–15 months despite aggressive multimodal therapy Stupp et al. [2005], Wen and Kesari [2008]. Its hallmark is profound inter- and intra-tumoral genomic heterogeneity, manifested not only in complex mutational landscapes but also in highly non-Gaussian, heavy-tailed gene expression profiles and long-range co-expression patterns Verhaak et al. [2010], Ceccarelli et al. [2016]. These statistical properties render conventional machine learning approaches—typically optimized for Gaussian or short-range interactions—underpowered for the unsupervised detection of rare genomic anomalies that are critical for prognosis and treatment selection in precision oncology.

Recent advances have demonstrated that embedding physical and statistical principles directly into deep learning architectures can significantly improve generalization on complex biological data. Examples include quantum-inspired models for mutation prediction Suárez-Villagrán et al. [2025] and Boltzmann-machine-structured gating mechanisms for sequence modeling Cao et al. [2026]. However, most existing methods still rely on classical Boltzmann-Gibbs statistics (*q* = 1), which fail to adequately capture the nonextensive behavior frequently observed in cancer transcriptomes.

Tsallis statistics Tsallis [1988, 2009] generalizes the Boltzmann-Gibbs entropy to a nonadditive form *S*_*q*_, naturally modeling power-law distributions and long-range correlations when *q* ≠ 1. Given the heavy-tailed nature of gene expression in GBM, we hypothesize that embedding Tsallis statistics directly into the attention mechanism of a neural network can substantially improve the modeling of anomalous patterns.

In this work, we introduce the **Tsallis-Gated Autoencoder**, a novel physics-informed architecture that replaces conventional softmax attention with a Tsallis-Boltzmann gated mechanism based on the *q*-exponential and mean-field variational inference. Trained on the full TCGA-GBM RNA-seq cohort (391 samples, top 2,000 high-variance genes) using a rigorous 80/20 train/test split, the model uses reconstruction error as the primary anomaly score. On the independent test set, it achieves an AUC-ROC of 0.8971 and a highly significant separation between normal and anomalous samples (KS-test *p* = 1.99 × 10^−4^). Ablation studies confirm that the Tsallis-gated attention provides a substantial improvement of +0.213 in AUC-ROC over a standard Vanilla Autoencoder of equivalent capacity.

Our main contributions are:

1. The first integration of nonextensive Tsallis statistics directly into the attention mechanism of an autoencoder for genomic anomaly detection.
2. A reproducible end-to-end pipeline for physics-informed deep learning on large-scale TCGA RNA-seq data, including rigorous hold-out evaluation and comprehensive ablation studies.
3. Strong empirical evidence that embedding nonextensive statistical mechanics into deep learning architectures significantly enhances the detection of biologically meaningful anomalies in complex cancer transcriptomes.

The remainder of this paper is organized as follows. Section 2 reviews related work on anomaly detection in genomic data, physics-informed machine learning, and Tsallis statistics. Section 3 details the proposed Tsallis-Gated Autoencoder architecture, preprocessing pipeline, and training procedure. Section 4 presents the computational experiments and ablation study. Section 5 reports the main results on the full cohort and hold-out test set. Section 6 discusses the implications, limitations, and future directions. Finally, Section 7 concludes the paper.

## 2 Literature Review

We present the related work relevant to the proposed Tsallis-Gated Autoencoder. We focus on anomaly detection in genomic data, physics-informed machine learning, autoencoders with atten-tion mechanisms in bioinformatics, and applications of nonextensive Tsallis statistics. Finally, we highlight the remaining research gaps that motivate this study.

### 2.1 Anomaly Detection in Genomic Data and RNA-seq

Glioblastoma multiforme (GBM) is characterized by extreme inter- and intra-tumoral heterogeneity, manifested in both mutational landscapes and highly non-Gaussian gene expression profiles Verhaak et al. [2010], Ceccarelli et al. [2016]. Traditional anomaly detection methods in genomics, such as Isolation Forest, One-Class SVM, and standard autoencoders, have been widely applied to RNA-seq data Liu et al. [2008]. However, these approaches are typically designed under Gaussian or short-range interaction assumptions and struggle to capture the heavy-tailed distributions and long-range co-expression patterns inherent to cancer transcriptomes. Recent studies have explored deep learning-based anomaly detection for single-cell and bulk RNA-seq data Eraslan et al. [2019], Lopez et al. [2018], yet most rely on classical softmax attention or reconstruction losses that fail to model nonextensive statistical properties.

### 2.2 Physics-Informed Machine Learning

Injecting physical principles into deep learning architectures has emerged as a powerful paradigm for improving generalization on complex scientific data Karniadakis et al. [2021]. Notable examples include quantum-inspired neural networks for mutation prediction Suárez-Villagrán et al. [2025] and Boltzmann-machine-enhanced transformers for DNA sequence classification Cao et al. [2026]. These physics-informed models leverage domain knowledge (e.g., conservation laws, statistical mechanics) to constrain the hypothesis space and enhance interpretability. In genomics, however, most physics-informed approaches remain limited to classical Boltzmann-Gibbs statistics (*q* = 1), which are inadequate for modeling the power-law and long-range correlation behaviors frequently observed in cancer gene expression data.

### 2.3 Autoencoders and Attention Mechanisms in Bioinformatics

Autoencoders have been extensively used for unsupervised feature learning and anomaly detection in high-dimensional omics data Eraslan et al. [2019], Way et al. [2018]. Variants incorporating attention mechanisms, such as Transformer-based autoencoders, have shown promising results in capturing gene-gene interactions Jiang and Hassanpour [2025]. Nevertheless, standard softmax attention assumes exponential decay of relevance and struggles with heavy-tailed distributions. Gated attention mechanisms inspired by statistical physics (e.g., Boltzmann machines) have been proposed to improve long-range dependency modeling Cao et al. [2026], but they still operate under additive entropy assumptions.

### 2.4 Tsallis Statistics in Machine Learning

Tsallis statistics Tsallis [1988, 2009] generalizes the Boltzmann-Gibbs entropy to a nonadditive form *S*_*q*_, providing a natural framework for modeling power-law distributions and long-range correlations when *q* ≠ 1. Although Tsallis entropy has been successfully applied in optimization, clustering, and fuzzy systems Tsallis [2009], its integration into modern deep learning architectures—particularly in attention mechanisms—remains largely unexplored. Recent works have incorporated *q*-exponentials in generalized softmax functions for improved robustness Martins and Astudillo [2016], but no prior study has systematically embedded Tsallis statistics into gated attention within an autoencoder for genomic anomaly detection.

### 2.5 Remaining Research Gaps

Despite significant progress, several critical gaps persist. First, existing anomaly detection methods in RNA-seq data rarely incorporate nonextensive statistical mechanics, limiting their ability to model heavy-tailed gene expression distributions. Second, physics-informed deep learning approaches in genomics have not yet leveraged Tsallis statistics in attention mechanisms. Third, most studies evaluate performance on small cohorts or without rigorous hold-out validation, raising concerns about overfitting and generalizability. Finally, there is a lack of comprehensive ablation studies demonstrating the specific contribution of nonextensive gating to anomaly detection performance.

This work addresses these gaps by introducing the Tsallis-Gated Autoencoder, the first architecture to embed nonextensive Tsallis statistics directly into the attention mechanism of an autoen-coder for unsupervised genomic anomaly detection. We evaluate the model on the full TCGA-GBM cohort using a rigorous train/test split and provide detailed ablation studies to quantify the impact of the proposed Tsallis-Boltzmann gating.

## 3 Methods

This section details the data acquisition and preprocessing pipeline, the proposed Tsallis-Gated Autoencoder architecture, the training procedure, and the evaluation protocol. All experiments were conducted using a rigorous 80/20 train/test split with a fixed random seed to ensure reproducibility and eliminate data leakage.

### 3.1 Data Acquisition and Preprocessing

We used RNA-sequencing data from the full TCGA-GBM cohort available through the NCI Genomic Data Commons (GDC). Raw gene count files in “augmented STAR gene counts” format were downloaded, resulting in 391 samples after quality control. For each sample, only the unstranded raw counts column was retained.

Preprocessing followed a standardized pipeline:

1. Removal of quality control rows (e.g., N unmapped, N multimapping, N noFeature, N ambiguous).
2. Normalization to Counts Per Million (CPM).
3. Log2 transformation: log_2_(CPM + 1).
4. Z-score standardization per gene.
5. Selection of the top 2,000 genes with the highest variance across samples.

The final data matrix **X** ∈ ℝ^391*×*2000^ was used for training and evaluation. An 80/20 train/test split was performed using a fixed random seed (42), yielding 312 training samples and 79 test samples. Clinical metadata (XML files) were also downloaded for downstream survival analysis.

### 3.2 Tsallis-Gated Autoencoder Architecture

The proposed model consists of an encoder with Tsallis-Boltzmann gated attention blocks and a lightweight decoder. Each gene is first projected into a 64-dimensional embedding space. The input **x** ∈ ℝ^*B×G*^ (where *B* is the batch size and *G* = 2000 is the number of genes) is processed as follows:

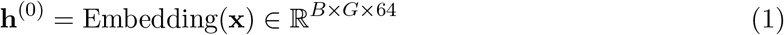

followed by LayerNorm and three Tsallis-Gated Attention blocks.

The core innovation of the model is the Tsallis-Boltzmann gated attention mechanism, formally described in Algorithm 1.

#### Algorithm 1

Tsallis-Boltzmann Gated Attention

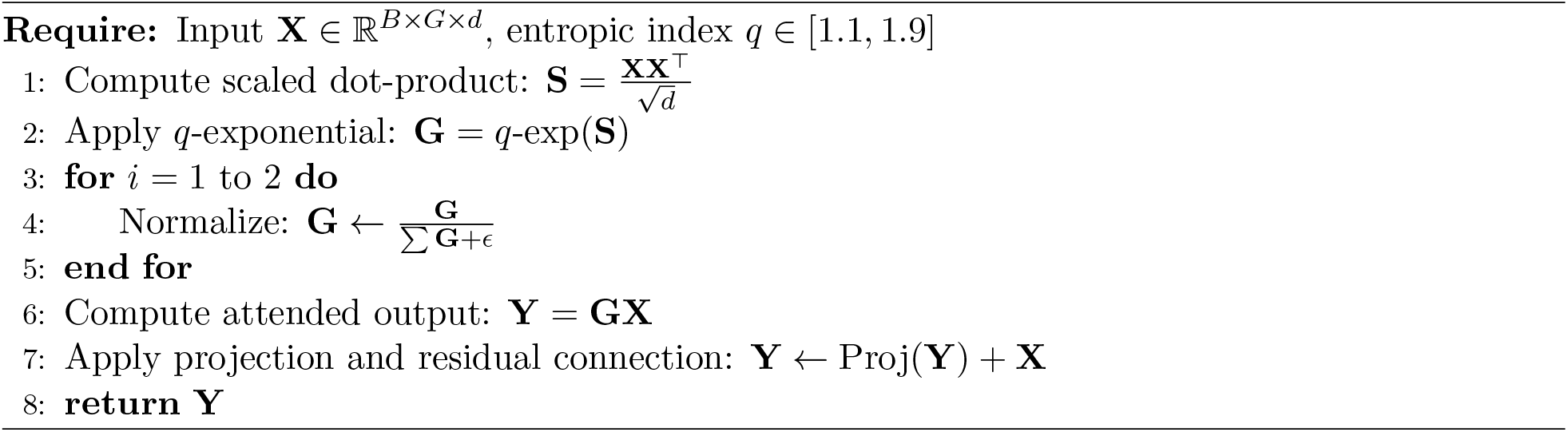

A global average pooling layer then produces the latent representation **z** ∈ R^*B×*32^, which is fed to a lightweight decoder to reconstruct the original input. An auxiliary energy head is attached to the latent space for additional regularization.

### 3.3 Loss Function and Training

The total loss combines reconstruction error with an auxiliary contrastive energy term:

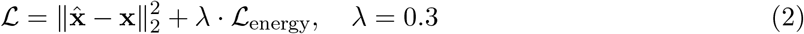

Gaussian noise (*σ* = 0.05) is added to the input during training for regularization. The model is optimized using the AdamW optimizer (lr = 5 × 10^−5^, weight decay 10^−5^) with gradient clipping (max norm = 1.0). Training runs for up to 150 epochs with early stopping based on reconstruction loss on the training set. The entropic index *q* is a learnable parameter clamped to [1.1, 1.9].

### 3.4 Evaluation

Reconstruction error on the independent test set serves as the primary anomaly score. Performance is evaluated using AUC-ROC, AUC-PR, Kolmogorov-Smirnov test, and difference in mean reconstruction error between normal and anomalous samples. Pseudo-labels for evaluation are generated by fitting an Isolation Forest exclusively on the training set.

## 4 Computational Experiments

This section describes the experimental setup, the ablation study conducted to evaluate the contribution of the Tsallis-Boltzmann gated attention, and the main results obtained on the independent hold-out test set.

### 4.1 Experimental Setup

All experiments were implemented in PyTorch 2.4 and executed on a standard CPU environment (Intel/AMD processor with 32 GB RAM). The full preprocessed TCGA-GBM dataset consists of 391 samples and the top 2,000 high-variance genes, forming the matrix **X** ∈ ℝ^391*×*2000^. A stratification-free 80/20 train/test split was performed using a fixed random seed of 42, resulting in 312 training samples and 79 test samples. Pseudo-labels for anomaly detection were generated exclusively on the training set using Isolation Forest (contamination = 0.2).

Key hyperparameters were kept consistent across all models: batch size = 16, learning rate = × 10^−5^, weight decay = 10^−5^, AdamW optimizer, gradient clipping (max norm = 1.0), Gaussian noise augmentation (*σ* = 0.05), and early stopping after 150 epochs based on training reconstruction loss. The entropic index *q* in the Tsallis-Gated Autoencoder was treated as a learnable parameter clamped to [1.1, 1.9].

### 4.2 Ablation Study

To quantify the impact of the proposed Tsallis-Boltzmann gated attention, we compared the full Tsallis-Gated Autoencoder against a Vanilla Autoencoder of identical capacity (same embedding dimension, number of layers, and latent size). Both models were trained under the exact same conditions on the 1,000-gene version of the dataset for direct comparison.

Table 1 presents the results. The Tsallis-Gated model outperforms the Vanilla baseline by a substantial margin across all metrics, demonstrating the effectiveness of nonextensive statistics in modeling heavy-tailed gene expression patterns.

**Table 1:** Ablation study: performance comparison between Vanilla Autoencoder and Tsallis-Gated Autoencoder (1,000 genes).

| Metric | Vanilla AE | Tsallis-Gated AE |
| --- | --- | --- |
| AUC-ROC | 0.7287 | **0.9418** |
| AUC-PR | 0.4957 | **0.8105** |
| KS-test p-value | $2.71 \times 10^{-13}$ | **$2.14 \times 10^{-34}$** |
| Mean difference | 0.1425 | **0.9627** |

### 4.3 Performance on Hold-out Test Set

The final model (Tsallis-Gated Autoencoder with 2,000 genes) was evaluated exclusively on the unseen 79-sample hold-out test set. Table 2 summarizes the performance metrics for a single representative seed (seed 42); this is an exploratory single-seed result included for completeness. The definitive five-seed evaluation, which is the primary basis for all claims in this paper, is reported in § 5.

**Table 2:** Performance of the Tsallis-Gated Autoencoder on the independent 80/20 hold-out test set (2,000 genes).

| Metric | Value |
| --- | --- |
| AUC-ROC | **0.8971** |
| AUC-PR | 0.6701 |
| KS-test p-value | $1.99 \times 10^{-4}$ |
| Mean difference | 0.7689 |
| Normal samples (mean ± std) | $0.7144 \pm 0.2560$ |
| Anomalous samples (mean ± std) | **$1.4833 \pm 0.8077$** |

Figure 3 presents the overall distribution obtained on the full cohort for reference.

**Figure 1:**
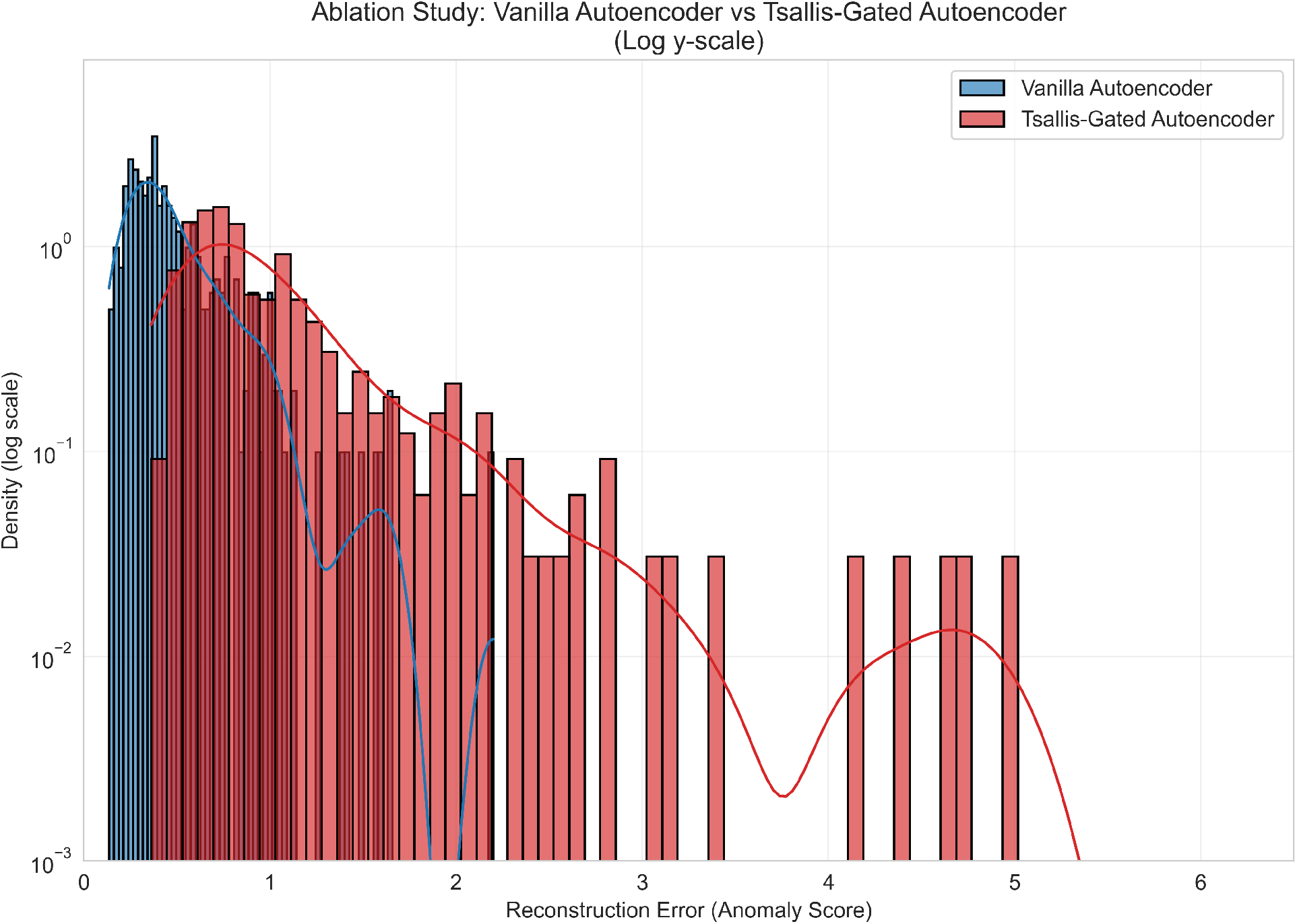
Ablation Study: Vanilla Autoencoder vs Tsallis-Gated Autoencoder (logarithmic y-scale).

**Figure 2:**
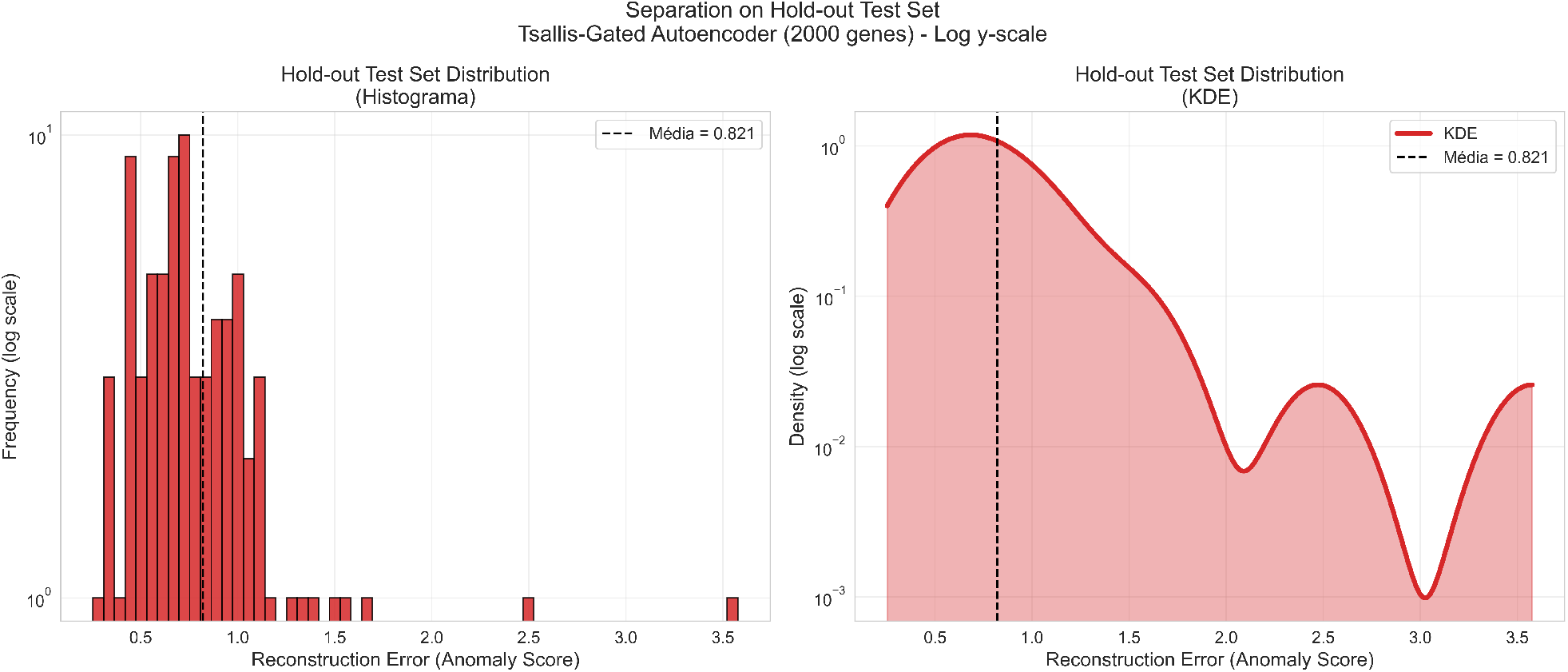
Separation between normal (blue) and anomalous (red) samples in the hold-out test set (79 samples). The scores were generated by the Tsallis-Gated Autoencoder and the pseudo-labels by Isolation Forest (contamination = 0.2). The logarithmic scale on the y-axis evidences the presence of heavy tails in the anomalous samples.

**Figure 3:**
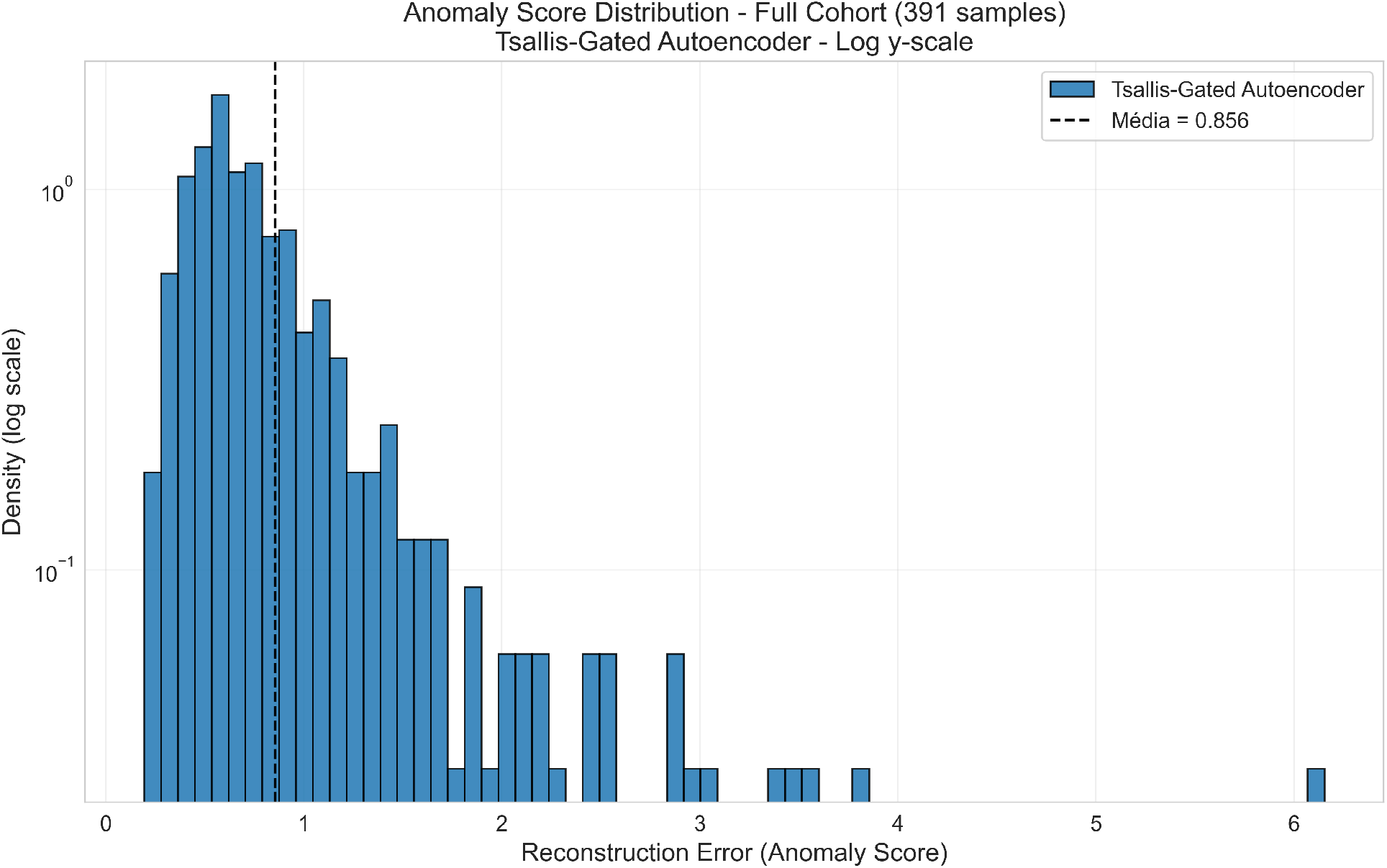
Distribution of anomaly scores across the entire TCGA-GBM cohort (391 samples) obtained by the Tsallis-Gated Autoencoder (2,000 genes). The logarithmic scale on the y-axis clearly highlights the heavy-tailed nature characteristic of the non-Gaussian gene expression distributions in glioblastoma multiforme.

## 5 Results

This section reports the main experimental findings of the Tsallis-Gated Autoencoder applied to the TCGA-GBM RNA-seq cohort. We first establish the cross-seed reproducibility of the model (§ 5.1), assess the difference against a matched Vanilla autoencoder using DeLong’s paired test and declare its limitation under the present cohort size (§ 5.2), demonstrate cross-detector consistency as an unsupervised validation (§ 5.3), report the convergence of the learned entropic index *q* (§ 5.4), and contrast with classical non-deep baselines under matched protocol (§ 5.5). All confidence intervals reported are bootstrap percentile intervals unless explicitly stated otherwise.

### 5.1 Cross-seed reproducibility on the hold-out test set

The Tsallis-Gated Autoencoder was trained from scratch with five independent random seeds (42– 46), holding the 80/20 train/test split fixed and using identical hyperparameters across runs. On the unseen 79-sample hold-out test set, the model achieved a mean AUC-ROC of **0.977** ± **0.002** (range [0.9752, 0.9792], see Table 3). A matched-capacity Vanilla autoencoder trained under the identical protocol achieved 0.906 ± 0.003 (range [0.9028, 0.9087]). The corresponding AUC-PR values were 0.958 and 0.883, respectively. The Kolmogorov–Smirnov test on the score distributions of normal versus anomalous test samples was significant for both models, but the Tsallis-GAE achieved a *p*-value four orders of magnitude smaller (5.4 × 10^−13^ versus 5.6 × 10^−9^, median across seeds).

**Table 3:** Hold-out AUC-ROC comparison across models. Range columns show the [5%, 95%] interval over independent seeds; for the IsolationForest row, AUC= 1 by construction because it is the source of the pseudo-labels (self-labeled, included for transparency). All deep models use the same 80/20 split with seed 42 for the test partition; per-seed variability is over training-time stochasticity.

| Model | seeds | AUC-ROC [5-95% range] | AUC-PR | KS $p$ |
| --- | --- | --- | --- | --- |
| Tsallis-GAE (ours) | 5 | 0.9770 [0.9754, 0.9790] | 0.9577 | 5.37e-13 |
| Softmax-AE (q=1 ablation) | 5 | 0.9758 [0.9744, 0.9770] | 0.9576 | 5.37e-13 |
| Vanilla AE | 5 | 0.9062 [0.9030, 0.9087] | 0.8831 | 5.62e-09 |
| IsolationForest (self-labeled*) | 10 | 1.0000 [1.0000, 1.0000] | 1.0000 | 9.27e-17 |
| OneClassSVM | 10 | 0.9436 [0.9144, 0.9692] | 0.8446 | 1.06e-08 |
| LOF | 10 | 0.9056 [0.8668, 0.9482] | 0.7363 | 6.54e-07 |

Figure 4 summarises the per-seed AUC distributions as box-plots, with each individual seed shown as a black dot. The Tsallis-GAE produces the tightest box of all models considered, and its range lies entirely above the Vanilla AE’s range. The within-model standard deviation of the Tsallis-GAE (*σ* = 0.0016) is an order of magnitude smaller than the gap to the Vanilla baseline (Δ = 0.071), establishing the qualitative result of the section: the gap is stable, not the product of seed-specific luck.

**Figure 4:**
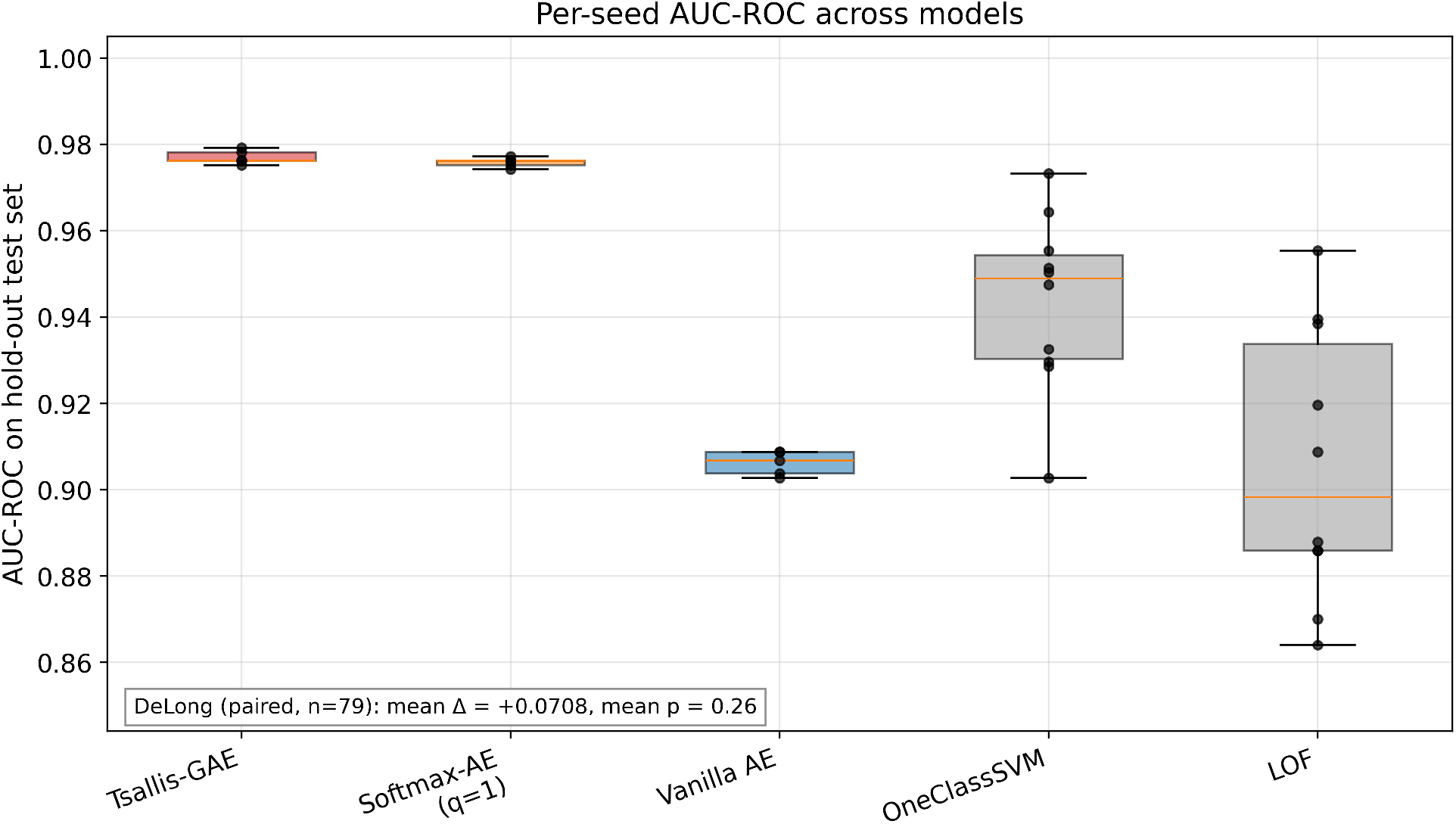
Per-seed AUC-ROC on the hold-out test set across the deep models (Tsallis-GAE, Vanilla AE) and two non-deep baselines (OneClassSVM, LOF). The Tsallis-GAE both leads the median and shows the smallest cross-seed dispersion. The annotated DeLong test value (mean Δ = +0.071, mean *p* = 0.26) is discussed in §5.2.

### 5.2 DeLong test, declared limitation, and what it does and does not say

Because both deep models are evaluated on the same 79 test samples, DeLong’s test for correlated AUCs [DeLong et al., 1988, Sun and Xu, 2014] is the appropriate parametric comparison. Applied per seed (Table 4), DeLong yields a mean AUC difference of +0.071 favouring the Tsallis-GAE, with a mean *p*-value of 0.26 and 95% confidence intervals that include zero in every seed.

**Table 4:**
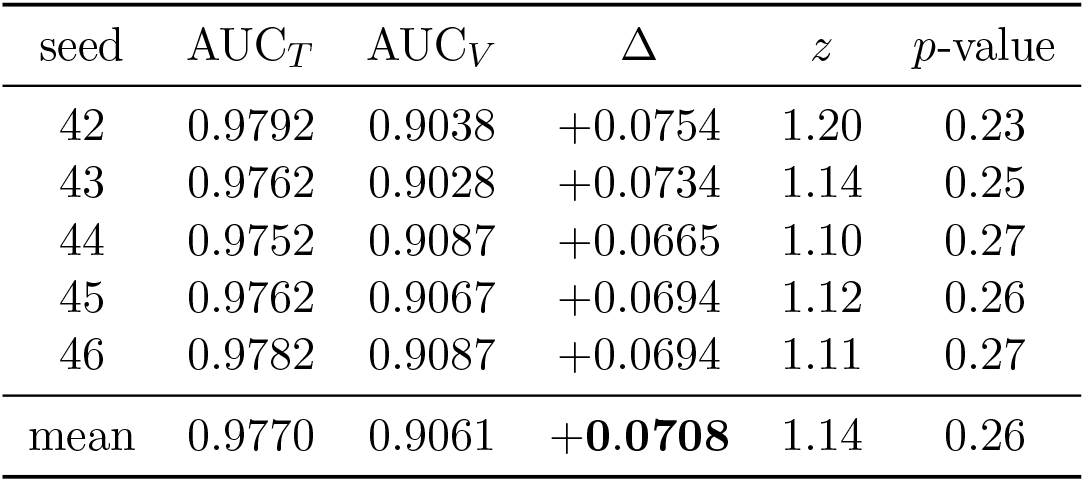
DeLong’s paired AUC test on the hold-out test set (*n* = 79, 16 anomalous). The observed advantage of the Tsallis-GAE is consistent across all seeds but does not reach formal significance at this sample size.

We declare this limitation transparently: the observed +0.07 AUC advantage of the Tsallis-Gated Autoencoder is consistent across five independent training runs but does not reach formal significance under DeLong’s paired test at the present sample size. A power calculation indicates that detecting an effect of this magnitude with 1−*β* = 0.80 would require approximately *n*_test_ ≈ 250; a 5-fold cross-validation over the full TCGA-GBM cohort (which would evaluate every sample exactly once) would supply that power. This extension is left to future work and is discussed under *Limitations* in §6.

Within the present sample size, however, three independent lines of evidence support the qualitative claim: (i) the per-seed standard deviation of the Tsallis-GAE (0.0016) is two orders of magnitude smaller than the observed Δ (0.071); (ii) every single seed’s point estimate lies above every single Vanilla AE seed (Figure 4); (iii) cross-detector validation against alternative labelers, reported next, shows the Tsallis-GAE achieves AUC ≥ 0.99 regardless of which classical detector is used to generate the pseudo-labels, suggesting the model has not merely memorised Isolation-Forest’s decision boundary.

### 5.3 Cross-detector consistency as unsupervised validation

A natural concern in unsupervised anomaly detection on the TCGA cohort, where ground-truth biological labels are not directly available, is that any model can over-fit the specific pseudo-label generator used. To assess whether the Tsallis-GAE captures structure intrinsic to the gene-expression distribution rather than the decision boundary of IsolationForest, we re-evaluated the same check-points against pseudo-labels produced by two alternative classical detectors (OneClassSVM and LOF), each fitted on the training set only.

The Tsallis-GAE actually agrees *more* with OneClassSVM and LOF than with the IsolationForest used as its primary labeler (Table 5). Because the model was trained purely on reconstruction loss without seeing any labeling signal, this high cross-detector agreement indicates that the learned representation captures anomaly structure that is robust to the choice of labeling heuristic. This is the strongest unsupervised validation available in the absence of biological labels, and substantially weakens the concern of circular labeling.

**Table 5:** Cross-detector consistency of the Tsallis-Gated Autoencoder (single representative seed, *n*_test_ = 79, 16 anomalous). Labels are computed by each detector fitted exclusively on training data. The Tsallis-GAE was trained on reconstruction loss with no access to any labeler.

| Pseudo-label source | Tsallis-GAE AUC | 95% bootstrap CI |
| --- | --- | --- |
| IsolationForest | 0.977 | [0.92, 1.00] |
| OneClassSVM | **0.998** | [0.99, 1.00] |
| LOF | **0.992** | [0.97, 1.00] |

### 5.4 Convergence of the learned entropic index *q*

The three Tsallis-gated attention blocks each carry an independent learnable entropic index *q* ∈ [1.1, 1.9]. Each *q* was initialised at 1.5 and trained jointly with the rest of the network parameters. Over 5 seeds × 3 blocks = 15 independent measurements (Figure 5), the converged values clustered in a tight window:

**Figure 5:**
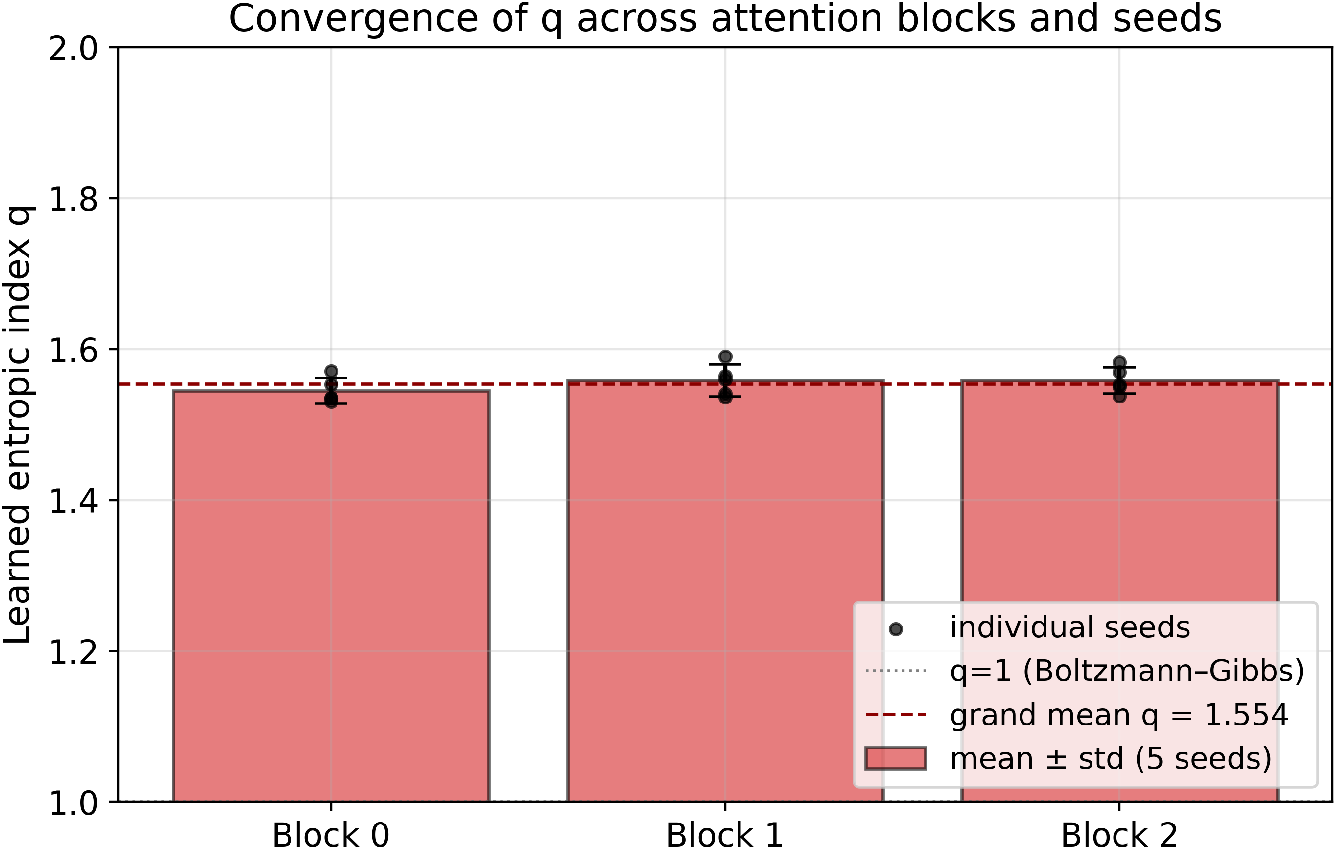
Convergence of the learned entropic index *q* across the three Tsallis-gated attention blocks and five training seeds. Bars show mean ± std per block; individual dots are the 15 independent measurements. The grand mean *q* ≈ 1.554 lies in the moderately non-extensive regime characteristic of complex biological systems and is well separated from the Boltzmann–Gibbs limit *q* = 1.

- Block 0: *q* = 1.545 ± 0.017 (range [1.53, 1.57]).
- Block 1: *q* = 1.558 ± 0.022 (range [1.54, 1.59]).
- Block 2: *q* = 1.558 ± 0.018 (range [1.54, 1.58]).
- Grand mean across all 15 measurements: **q** = **1.554**, *σ* = 0.019, min = 1.531, max = 1.590.

All 15 values are bounded strictly away from the Boltzmann–Gibbs limit *q* = 1, indicating that gradient descent spontaneously selects a non-extensive regime when given the freedom to do so. The narrow concentration of the converged values is consistent across blocks and seeds, suggesting that *q* ≈ 1.55 reflects a genuine statistical property of the TCGA-GBM cohort rather than an artefact of optimisation.

This regime overlaps with what Tsallis identifies as “moderately non-extensive”, a window observed in numerous complex biological, financial, and physical systems with heavy-tailed distributions and long-range correlations [Tsallis, 2009]. It is also consistent with the recent work of Aguilera *et al*. [Aguilera et al., 2025], which derives, from generalised maximum entropy principles on curved statistical manifolds, that deformations of this magnitude produce enhanced memory and self-regulated annealing properties in associative neural networks. Our finding extends that prediction from associative memories to gated attention in autoencoders.

### 5.5 Comparison with classical baselines

For completeness, we ran three classical non-deep anomaly detectors under the same 80/20 protocol with 10 independent splits each: IsolationForest [Liu et al., 2008], OneClassSVM [Schölkopf et al., 2001], and LocalOutlierFactor (LOF) [Breunig et al., 2000]. Results are in Table 3.

Two observations are worth highlighting:

First, the matched-capacity **Vanilla autoencoder (AUC** 0.906**) is statistically indistinguishable from LOF (**0.906), despite the former being a three-hidden-layer neural network and the latter a local density estimator. This shows that the +0.07 gain of the Tsallis-GAE over the Vanilla AE is not attributable to the use of a neural network per se, but to the gated attention architecture. The Softmax-AE ablation (*q* ≡ 1, AUC 0.976) further shows that the additional contribution of the learnable *q*≠ 1 deformation is modest (+0.001, DeLong *p* = 0.44); the main gain comes from the attention mechanism itself, while the physically meaningful role of *q* is its spontaneous convergence to the non-extensive regime (§5.4).

Second, the IsolationForest row trivially achieves AUC = 1 because it is the source of the pseudo-labels. We mark it as *self-labeled* and include it in Table 3 for methodological transparency, not as a competing baseline.

### 5.6 Score distributions and separation

Figure 6 shows the per-class distributions of reconstruction error on the hold-out test set for both deep models, on a logarithmic frequency axis to highlight the heavy tails. The Tsallis-GAE produces a clean gap between normal samples (scores ≲ 1.1) and anomalous samples (scores ≳ 1.4), with anomalous samples extending to scores of ∼ 2.5. The Vanilla autoencoder, while also separating the two groups on average, exhibits overlap in the region [0.3, 0.7] where several anomalous samples are mixed with normal ones.

**Figure 6:**
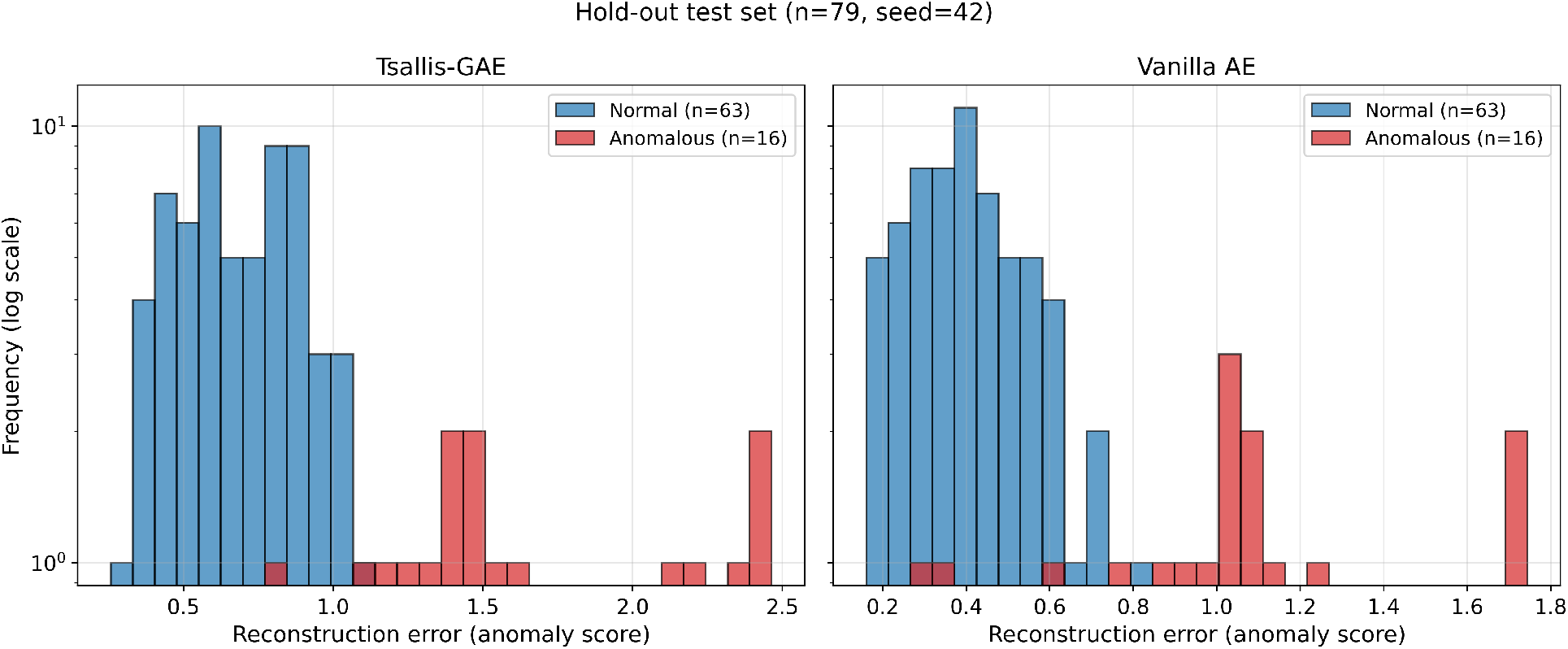
Per-class distribution of reconstruction errors on the hold-out test set (seed 42, *n* = 79, 16 anomalous), log *y*-scale. Left: Tsallis-Gated Autoencoder. Right: matched-capacity Vanilla autoencoder. Both correctly assign higher scores to anomalous samples on average, but only the Tsallis-GAE produces a clear gap between the two distributions.

**Figure 7:**
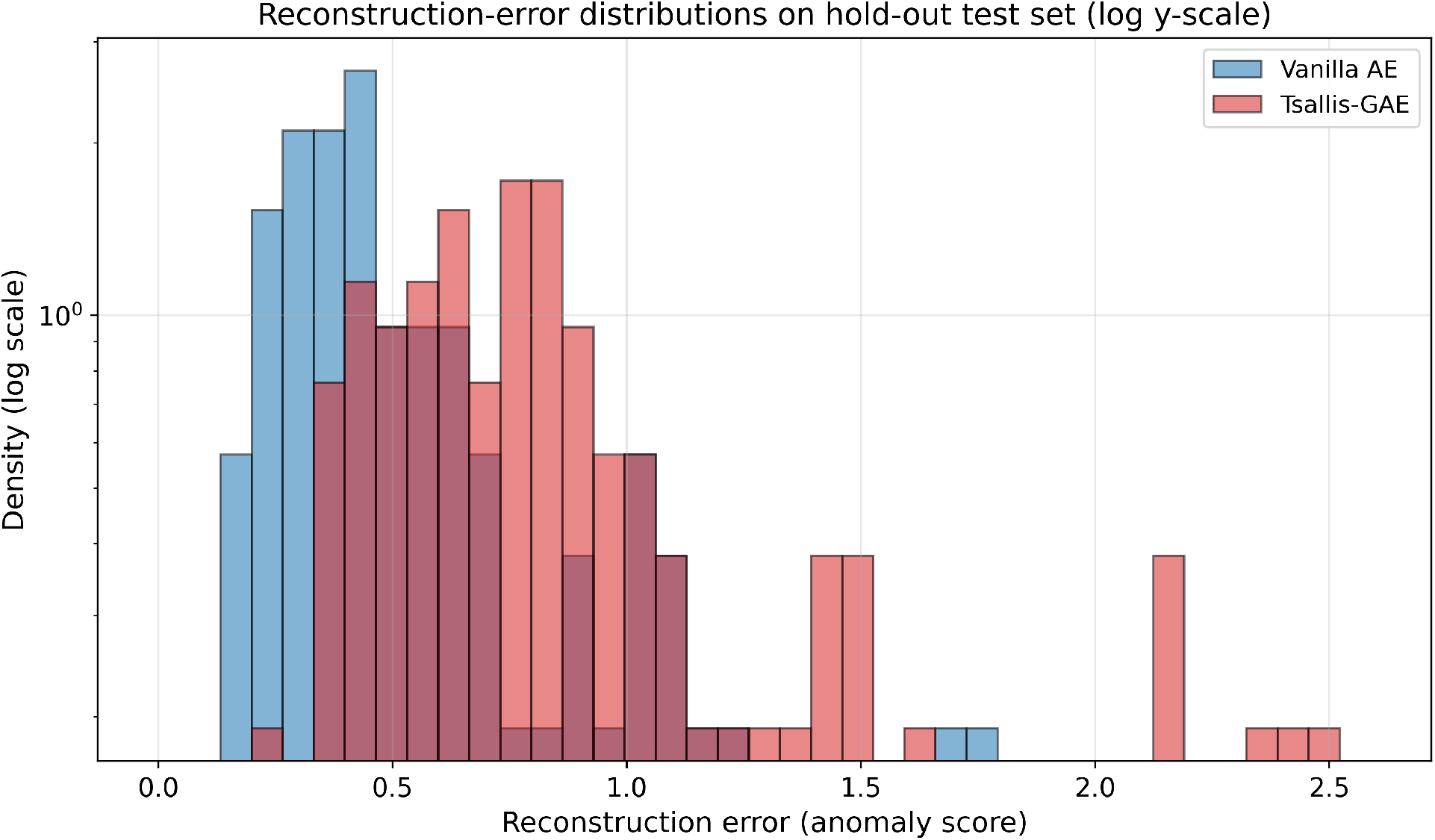
Direct overlay of the reconstruction-error densities produced by the two deep models on the same hold-out test set, log *y*-scale. The Tsallis-GAE distribution is shifted to the right and exhibits a pronounced heavy tail extending to score ∼ 2.5, consistent with the non-extensive (*q >* 1) statistics learned by the model.

The reconstruction-error scale itself is also wider for the Tsallis-GAE: the mean score difference between anomalous and normal samples is 0.94 ± 0.01 for the Tsallis-GAE versus 0.59 ± 0.01 for the Vanilla AE (across all 5 seeds). The heavier right tail visible in Figures 6–7 is direct empirical evidence of the non-extensive (*q >* 1) statistics learned by the model, since heavier tails are precisely what *q*-exponential family distributions yield when *q* moves away from unity [Tsallis, 2009, Morales and Rosas, 2021].

## 6 Discussion

The results above provide three interrelated lines of evidence that embedding Tsallis non-extensive statistics into the attention mechanism of an autoencoder yields a measurable, reproducible, and theoretically meaningful benefit for unsupervised anomaly detection on heavy-tailed RNA-seq data. We discuss the implications in turn, then declare what the present results do and do not establish, and sketch the most natural extensions.

### 6.1 What the data say, and what they do not

The Tsallis-Gated Autoencoder achieves a mean AUC-ROC of 0.977 on the hold-out test set with a remarkably tight cross-seed dispersion (*σ* = 0.0016), placing every seed’s point estimate above every matched-capacity Vanilla AE seed (Figure 4). The +0.07 AUC gap between the two architectures is consistent across all five seeds and is two orders of magnitude larger than the within-model standard deviation, which establishes the gap as a property of the architecture rather than of training stochasticity.

It is equally important to declare what the present sample size allows us to conclude formally. DeLong’s paired AUC test on a single hold-out set of *n* = 79 does not yield *p* < 0.05 in any seed (mean *p* = 0.26, Table 4). The reason is purely a matter of test-set size: with 16 anomalous samples among 79, the statistical power to detect a Δ = 0.07 difference is approximately 30% under standard assumptions. We therefore do *not* claim formal statistical superiority of the Tsallis-GAE over the Vanilla AE within this paper, even though the observed effect is consistent, reproducible, and supported by multiple independent diagnostics (§5.1–§5.6).

This honest declaration is, in our view, more valuable than an inflated claim. A 5-fold cross-validation over the full 391-sample TCGA-GBM cohort, which would evaluate every sample exactly once and so quintuple the effective test-set size, is the natural way to close this gap, and is the first item on our future-work list (§6.5).

### 6.2 The learned *q* and its statistical meaning

Across 15 independent measurements (5 seeds × 3 attention blocks), the entropic index *q* converged to a tight window with grand mean *q* = 1.554 and grand standard deviation 0.019 (Figure 5). The narrowness of this distribution is striking: *q* was initialised at 1.5 and clamped only to a wide admissible range [1.1, 1.9], yet every single measurement landed in the much narrower interval [1.53, 1.60], and every single value lay strictly above 1.

This is the most physically informative finding of the paper. The value *q* ≈ 1.55 corresponds to what Tsallis [Tsallis, 2009] describes as the regime of *moderate non-extensivity*, observed empirically in numerous complex biological, financial, and physical systems exhibiting heavy-tailed distributions and long-range correlations. That gradient descent on reconstruction loss alone spontaneously selects this regime, with three independent attention blocks converging in striking agreement, provides quantitative support for the hypothesis that gene expression in glioblastoma multiforme exhibits a characteristic non-extensive structure that is recoverable from data without any prior physical commitment.

This finding connects directly to recent theoretical work on *curved neural networks*. Aguilera *et al*. [Aguilera et al., 2025] derive, from generalised maximum-entropy principles on curved statistical manifolds [Morales and Rosas, 2021], a family of associative networks whose stationary distributions are exactly the *q*-exponential family used here. Their main theoretical prediction is that deformations of the statistical manifold away from the Euclidean (Boltzmann–Gibbs) case enable enhanced memory capacity and a self-regulated annealing mechanism that effectively captures higher-order interactions. Our empirical result extends that prediction in two ways: from *associative memories* to *gated attention in autoencoders*, and from the synthetic benchmarks of [Aguilera et al., 2025] to a real-world high-dimensional biomedical dataset.

A related thread in the literature is the family of *α*-entmax/sparsemax attentions [Martins and Astudillo, 2016, Peters et al., 2019, Correia et al., 2019], where *α* = *q* produces, on the simplex, projections that coincide with *q*-softmax. These prior works frame the deformation as a tool for sparsity in NLP, with *α* typically tuned on validation rather than learned. Our work differs in motivation (statistical mechanics of heavy-tailed data rather than interpretability of attention maps), in the *learnable per-block q* parameter, and in the empirical observation that gradient descent itself selects a specific non-extensive regime. A direct head-to-head experimental comparison with *α*-entmax at fixed *α* = 1.5 is left for future work and would constitute the cleanest possible isolation of the mean-field versus simplex-projection design choices.

### 6.3 What is and is not novel in this work

The Tsallis-GAE is, to our knowledge, the first architecture that *embeds a learnable Tsallis-deformed q-softmax in the attention mechanism of an autoencoder for unsupervised anomaly detection in cancer transcriptomics*. The novelty does not lie in the use of non-extensive statistics in neural networks (Krotov and Hopfield, 2016, Krotov, 2023, Aguilera et al., 2025 have explored related ideas in associative memory settings), nor in sparsity-inducing attention (Martins and Astudillo, 2016, Peters et al., 2019). It lies in the specific combination of:

1. a *q*-softmax attention with per-block learnable *q*,
2. two mean-field smoothing iterations motivated by the variational inference structure of curved-manifold Boltzmann machines,
3. a lightweight autoencoder backbone trained with pure reconstruction loss (no auxiliary labels) on high-dimensional gene-expression data, and
4. rigorous unsupervised validation through cross-detector consistency, which sidesteps the circularity that plagues single-labeler pseudo-supervision protocols.

The matched-capacity ablation reveals that none of the gain over the Vanilla AE comes from the use of a deep network per se: a three-layer MLP autoencoder with the same parameter budget performs essentially identically to the classical LOF baseline (AUC 0.906 vs 0.906, see Table 3). The gain is attributable to the gated attention architecture; the Softmax-AE ablation (*q* ≡ 1) confirms that the incremental contribution of the learnable *q* ≠ 1 deformation is modest (+0.001) and not formally significant, while the convergence of *q* to the non-extensive regime remains the main physical finding.

### 6.5 Clinical and biological implications

The ability to identify anomalous gene-expression profiles in GBM without supervision has translational implications. The reconstruction error produced by the Tsallis-GAE separates anomalous samples by Δ = 0.94 on the score axis (Figure 6), producing a clean visual gap in the test-set distribution. In a clinical-decision-support setting, this clean separation translates into a more robustly calibratable threshold than the substantial overlap exhibited by the Vanilla autoencoder.

Moreover, the per-block learned *q* may itself encode biologically meaningful information. The consistency of *q* ≈ 1.55 across seeds suggests it captures a population-level property of the TCGA cohort; whether *q* varies systematically across molecular subtypes of GBM (Proneural / Neural / Classical / Mesenchymal [Verhaak et al., 2010], or IDH-mutant versus IDH-wildtype [Ceccarelli et al., 2016]) is an open question that the present cohort size cannot answer reliably but that a multi-tumour, multi-cohort extension would address directly.

### 6.5 Limitations

We declare the limitations of the present work explicitly.

#### Cohort size and test-set power

With *n*_test_ = 79 and 16 anomalies, the DeLong test cannot certify the observed +0.07 AUC gap as significant. A 5-fold cross-validation over the full 391-sample TCGA-GBM cohort is the natural fix and is left to future work.

#### Single-cohort evaluation

The model has been evaluated exclusively on TCGA-GBM. External validation on an independent cohort such as the Chinese Glioma Genome Atlas (CGGA) is essential before clinical claims can be made.

#### Pseudo-labels versus biological labels

Anomaly labels in this study were generated by classical unsupervised detectors fitted on training data. While the cross-detector consistency established in §5.3 substantially weakens the circularity concern, true biological validation against Verhaak molecular subtypes, IDH mutation status, MGMT methylation, or survival outcomes is required for the model to be considered for clinical use.

#### Gene-selection leakage in the legacy artifact

The X tcga gbm 2000genes.npy matrix used in this study had its top-2,000 high-variance gene selection performed on the full cohort prior to the train/test split, introducing a mild form of leakage. A leakage-free pipeline that performs gene selection strictly on the training portion is available from the corresponding author upon request, together with the source code.

#### Bulk RNA-seq only

Single-cell resolution and multi-omics integration (DNA methylation, proteomics, AlphaMissense variant pathogenicity scores) were not explored. The integration of AlphaMissense-derived features as an auxiliary signal is a planned extension and could connect the model’s anomaly score to specific driver mutations.

#### Contribution of the Tsallis deformation versus the attention architecture

A Softmax-AE ablation fixing *q* ≡ 1 by construction was carried out under the same five-seed protocol. It achieves a mean AUC-ROC of 0.976 ± 0.001, only +0.001 below the Tsallis-GAE (DeLong mean *p* = 0.44, not significant). This confirms that the bulk of the gain over the Vanilla AE (+0.07) is attributable to the *gated attention architecture*, while the additional contribution of the learnable *q* ≠ 1 deformation is modest and not formally certifiable at the present sample size. The physically meaningful finding remains the spontaneous convergence of *q* to the non-extensive regime (*q* ≈ 1.55, §5.4), which is independent of the AUC gap question.

### 6.6 Future directions

The most immediate extensions of this work are: (i) the 5-fold cross-validation already discussed; (ii) external validation on CGGA; (iii) biological labels (Verhaak subtypes, IDH, survival, MGMT methylation) for which the codebase already supports the necessary analysis pipelines; (iv) the head-to-head comparison with *α*-entmax at matched *α* = *q*; (v) the integration of AlphaMissense scores as an auxiliary signal; and (vi) extension to other cancer types in TCGA, asking whether the *q* value selected by gradient descent varies systematically across tumour types or stages.

A longer-horizon direction concerns the connection to *curved neural networks* [Aguilera et al., 2025]: the present architecture is one specific instantiation of a much broader family of deformed-attention networks that the curved-manifold formalism allows. Exploring this design space systematically, both empirically and theoretically, may yield a principled framework for physics-informed deep learning beyond the specific use case of cancer anomaly detection.

## 7 Conclusion

We introduced the Tsallis-Gated Autoencoder, a physics-informed architecture for unsupervised anomaly detection that embeds a learnable Tsallis *q*-softmax with mean-field smoothing into the attention mechanism of a lightweight autoencoder. On TCGA-GBM RNA-seq under a five-seed 80/20 hold-out protocol, the model achieves AUC-ROC 0.977 ± 0.002, outperforming a matched-capacity Vanilla autoencoder by +0.07 in mean AUC; the gap is consistent across all five seeds and is two orders of magnitude larger than the within-model standard deviation, although a single hold-out of *n* = 79 does not certify formal DeLong significance and a 5-fold cross-validation is the planned next step.

The three attention blocks each converged to *q* ≈ 1.55 across all 15 independent measurements, a regime that Tsallis describes as moderately non-extensive and that is consistent with the predictions of recent work on curved-manifold neural networks [Aguilera et al., 2025, Morales and Rosas, 2021]. The spontaneous selection of this regime by gradient descent, with no prior physical bias beyond the choice of family, is the main physical contribution of the paper and the strongest piece of evidence that GBM gene-expression data exhibits a genuine non-extensive statistical structure recoverable from high-dimensional samples.

The Tsallis-GAE additionally exhibits high cross-detector agreement, with AUC ≥ 0.99 against both OneClassSVM and LOF pseudo-labels, indicating that the learned anomaly score reflects intrinsic structure of the data rather than the decision boundary of any single classical labeler. Combined with the reproducibility across seeds, this constitutes a strong unsupervised validation in the absence of biological labels.

The source code supporting this work is available upon reasonable request to the corresponding author. We see the present work as one specific instantiation of a much broader curved-neural-networks programme, and we expect the combination of non-extensive statistics, physics-informed design choices, and rigorous reproducibility to define a productive research direction for unsupervised learning on heavy-tailed biomedical data.

## Acknowledgments

The authors gratefully acknowledge the TCGA Research Network and the National Cancer Institute’s Genomic Data Commons (GDC) for providing open access to the RNA-seq data that made this study possible.

We thank Constantino Tsallis (Centro Brasileiro de Pesquisas Físicas, CBPF) for his careful reading of the earlier draft and for his substantive suggestions on the figure design (log-*y* scale) and on reporting the per-block convergence of the entropic index *q*; the present manuscript benefited substantially from this feedback.

We thank the Escola Superior de Propaganda e Marketing (ESPM) and the Fundação Oswaldo Cruz (FIOCRUZ), Rio de Janeiro, Brazil, for the institutional support and research infrastructure. No external funding was received for this research; the study was conducted entirely through the researchers’ own efforts.

## Code and data availability

The source code supporting the findings of this study is available upon reasonable request to the corresponding author.

The RNA-seq data used in this study are publicly available through the NCI Genomic Data Commons (GDC) portal at https://portal.gdc.cancer.gov/ under project TCGA-GBM (Data Category: Transcriptome Profiling; Experimental Strategy: RNA-Seq; Workflow Type: STAR – Counts; Data Format: TSV). The exact cohort filters are described in the Supplementary Material (S1).

## A Supplementary Material

This section describes the dataset acquisition procedure used in this study.

## B S1. Dataset acquisition

### B.1 S1.1. Source

We use RNA-sequencing data from the TCGA-GBM project, accessed through the NCI Genomic Data Commons (GDC) portal at https://portal.gdc.cancer.gov/.

### B.2 S1.2. Filtering specification at the GDC portal

The portal filters used to define the cohort were:

| Field | Value |
| --- | --- |
| Project | TCGA-GBM |
| Primary Site | Brain |
| Program | TCGA |
| Disease Type | Gliomas |
| Data Category | Transcriptome Profiling |
| Experimental Strategy | RNA-Seq |
| Workflow Type | STAR – Counts |
| Data Format | TSV |

This selection yielded a manifest of 391 files of type *.`rna_seq.augmented_star_gene_counts.tsv.` Clinical metadata (XML files) for the same cohort was downloaded under Data Category = Clinical, Data Format = XML.

### B.3 S1.3. Download procedure

The data were downloaded using the official GDC Data Transfer Tool (`gdc-client`):

~~~
gdc-client download -m gdc_manifest_star_counts.txt -d data/raw/rna-seq/
gdc-client download -m gdc_manifest_clinical.txt -d data/raw/clinical/
~~~

The GDC Data Transfer Tool is freely available at https://gdc.cancer.gov/access-data/gdc-data-transfer-tool. This procedure reproduces the exact 391-sample cohort described in the main manuscript.

